# MoSBi: Automated signature mining for molecular stratification and subtyping

**DOI:** 10.1101/2021.09.30.462567

**Authors:** Tim Daniel Rose, Thibault Bechtler, Octavia-Andreea Ciora, Kim Anh Lilian Le, Florian Molnar, Nikolai Koehler, Jan Baumbach, Richard Röttger, Josch Konstantin Pauling

## Abstract

The improving access to increasing amounts of biomedical data provides completely new chances for advanced patient stratification and disease subtyping strategies. This requires computational tools that produce uniformly robust results across highly heterogeneous molecular data. Unsupervised machine learning methodologies are able to discover de-novo patterns in such data. Biclustering is especially suited by simultaneously identifying sample groups and corresponding feature sets across heterogeneous omics data. The performance of available biclustering algorithms heavily depends on individual parameterization and varies with their application. Here, we developed MoSBi (Molecular Signature identification using Biclustering), an automated multi-algorithm ensemble approach that integrates results utilizing an error model-supported similarity network. We evaluated the performance of MoSBi on transcriptomics, proteomics and metabolomics data, as well as synthetic datasets covering various data properties. Profiting from multi-algorithm integration, MoSBi identified robust group and disease specific signatures across all scenarios overcoming single algorithm specificities. Furthermore, we developed a scalable network-based visualization of bicluster communities that support biological hypothesis generation. MoSBi is available as an R package and web-service to make automated biclustering analysis accessible for application in molecular sample stratification.

## Introduction

Optimizing treatments and improving patients’ health is the goal of precision medicine. In contrast to canonical medicine, where treatments are prescribed empirically^1^, precision medicine aims to identify individually adapted treatments. Nowadays, diseases are commonly diagnosed based on the international classification of diseases (ICD). This assumes that a disease shows similar symptoms in every individual, hence treatments are meant to act on the majority of symptoms. Patient stratification for precision medicine builds on the idea that a cohort of patients with varying or similar symptoms might have different molecular causes. They can then be stratified on the molecular level and divided into subgroups^2^. Therefore, precision medicine wants to move away from classical disease definitions to characteristic signatures of molecular alterations which enable individualized treatments.

Achieving this requires understanding of molecular disease mechanisms. Unsupervised machine learning methods are best-suited, since they uncover the inherent structure of the given data and do not require labeled data, which might be biased towards classical disease understandings^3^. Unsupervised clustering methods seek to identify distinct subgroups over the entire features set; but it is unrealistic to assume that diseases manifest in all features. Instead they are limited to a subtype specific subset. Biclustering algorithms can meet this requirement.

Molecular data is usually available in data matrices with patient samples as columns and biomolecular features as rows. Biclustering algorithms cluster samples and biomolecules of a data matrix simultaneously. This results in sample groups with a molecular subset which characterizes the group. Numerous algorithms have been published, which try to tackle the problem from different angles. An overview about important concepts was published by Madeira and Oliveira^4^.

Similar to clustering^5^, evaluations of biclustering algorithms have shown differences in the performance under various real-life and synthetic conditions^6,7^. A common way to improve the results of machine learning techniques are ensemble approaches. The goal is to improve robustness, consistency, novelty and stability over what single algorithms could achieve^8^. Also for biclustering problems ensemble algorithms have been proposed^9–15^. Most of these ideas are adaptations of approaches for ensemble clustering. Some of the proposed methods are not implemented^9,12^ and therefore not easily accessible, while others are single algorithm ensemble approaches which cannot overcome limitations of one algorithm.

The analysis and interpretation of biclustering results can profit from visualizations, which show the content or relations between biclusters. Many approaches have been developed^16–22^, which are often bound to specific algorithms or do not scale well for many biclusters^17^.

Here we propose a multi-algorithm biclustering ensemble approach for the stratification of molecular samples. In the manuscript, we 1) Introduce the methodology and network visualization. 2) Evaluate the performance on multiple experimental metabolomics, proteomics and transcriptomics datasets. 3) With a framework for synthetic data generation, evaluate the approach on synthetic data. 4) Apply our approach in a multi-omics context and 5) Present open source software to make bicustering more accessible for research.

## Results

### A multi-algorithm ensemble biclustering approach

The steps of our ensemble approach (MoSBi - Molecular signature identification using biclustering) are described in **Figure 1A**, for full details, please refer to the methods section. At first, we selected a set of established or recently developed biclustering algorithms (**Table 1**), which are executed independently. Next, similarities between all biclusters are calculated. The similarity is described by the degree of overlap, meaning, the more samples and features shared between biclusters, results in higher similarity. Highly similar biclusters point towards the same pattern in the data. Similarities are filtered for random overlaps and a bicluster network is generated with biclusters as nodes and connections between them if they exceed a higher than random similarity. The example network shown in Figure 1A reveals several highly connected communities in the network, which are not as strongly connected between each other. By using the Louvain modularity, such communities can be extracted and converted into ensemble biclusters. Two thresholds control the size of resulting ensemble biclusters. We previously successfully utilized the principle of MoSBi to identify de-novo subtypes of non-alcoholic liver disease based on clinical lipidomics data^23^.

**Figure 1:**
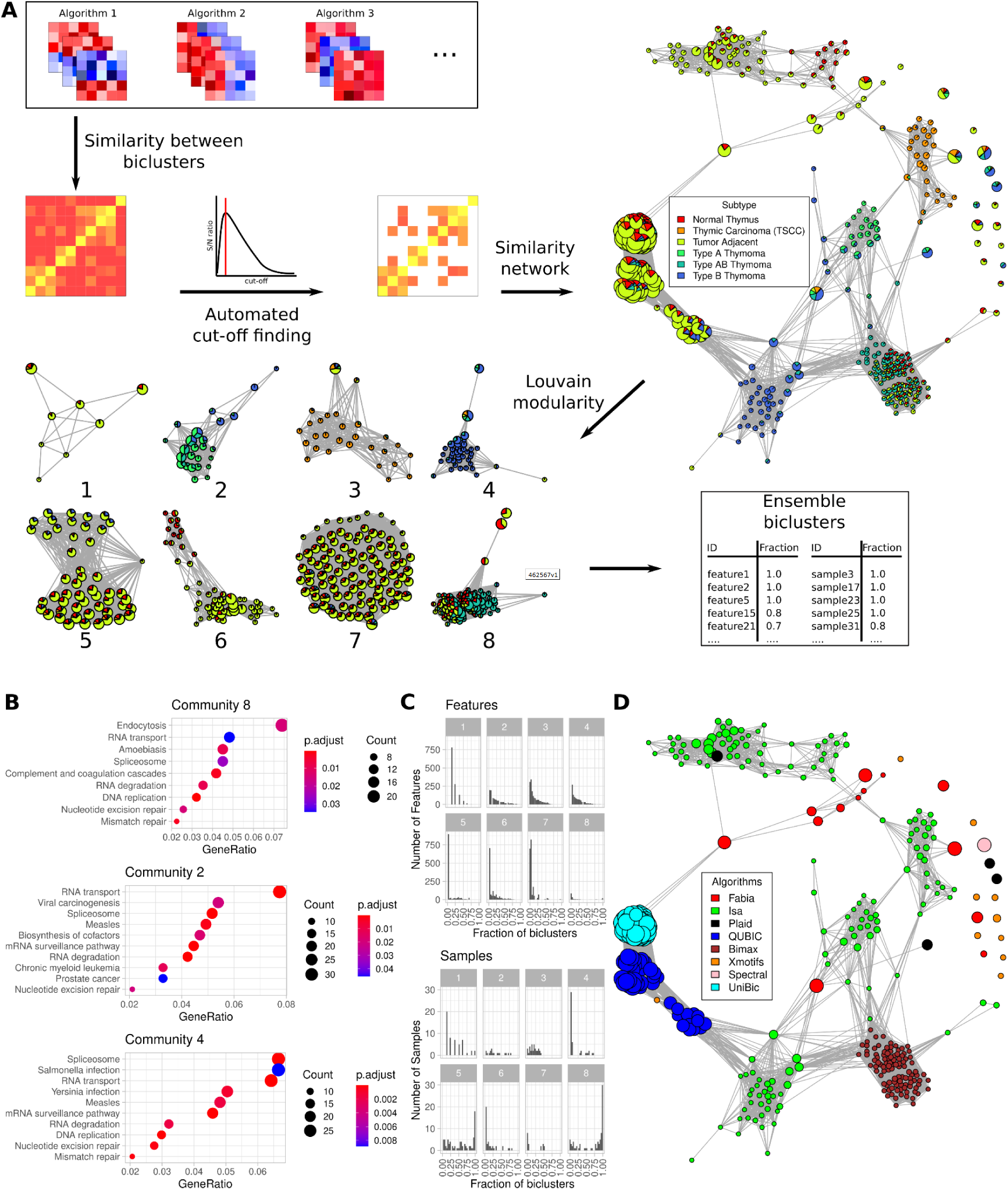
Workflow of MoSBi with exemplary network visualizations. (**A**) Steps of the MoSBi approach. First, biclusters are predicted by multiple algorithms and a similarity matrix is computed, which is then filtered for larger than random overlaps using an error model. The matrix is then converted to a network which can be visualized with meta-information about samples or features. Louvain communities are then extracted and converted into ensemble biclusters. As an example the bicluster network of proteomics data from Ku et al.^34^ is shown. Nodes represent biclusters, with edges between them if their overlap exceeds the error threshold. (**B**) KEGG pathway enrichment for features of selected communities 2, 4 & 8. (**C**) Frequency of features (upper) and samples (lower) in biclusters that belong to one community. (**D**) Bicluster network of proteomics data from Ku et al.^34^. Node colors represent algorithms, by which they were predicted.

**Table 1:** List of evaluated biclustering algorithms in alphabetical order. The results of algorithms can be imported and accessed with our MoSBi R package or executed using the webtool.

| Algorithm | Publication |
| --- | --- |
| BicARE | Gestraud et al. (2020) <sup>24</sup> |
| Bimax | Prelić et al. (2006) <sup>25</sup> |
| CC | Cheng & Church (2000) <sup>26</sup> |
| Fabia | Hochreiter et al. (2010) <sup>27</sup> |
| Isa | Bergmann et al. (2003) <sup>28</sup> |
| Plaid | Lazzeroni and Owen (2002) <sup>29</sup> |
| QUBIC | Zhang et al. (2010) <sup>30</sup> |
| Quest | Murali et al. (2002) <sup>31</sup> |
| Spectral | Kluger et al. (2003) <sup>32</sup> |
| UniBic | Wang et al. (2016) <sup>33</sup> |
| Xmotifs | Murali et al. (2002) <sup>31</sup> |

Before evaluating the performance of MoSBi on multiple omics datasets, we selected a public thymic epithelial tumor dataset^34^ to show the application and potential of our approach. Ku et al.^34^ measured the proteome of 134 tumor, tumor adjacent and normal thymus samples and revealed significant differences in the proteome signatures of thymoma subtypes. In **Figure 1A** the similarity network of predicted biclusters colored by sample groups can be seen. Additionally, node sizes were scaled according to the number of samples. It provides an overview of the match of predicted biclusters with known information about samples, in this case cancer subtypes/tissues. While being a central part of the workflow, to compute ensemble biclusters, networks also serve as a visualization of biclustering predictions.

It is immediately obvious that clusters of nodes can be found in the network, indicating biclusters with a similar set of samples and features. This can be observed by similar color distributions of biclusters clustering together. After applying the Louvain modularity, these clusters result in network communities.

Some communities, in particular 2, 4 & 8 predominantly consist of type A, B and AB thymoma. We performed KEGG pathway enrichment of protein sets from the ensemble biclusters (**Figure 1B**). All selected communities showed significant repair mechanism pathways, which is well known for tumors to influence those pathways. Additionally community 2, which includes samples of all thymoma subtypes, indicating a common signature on the proteomic level, showed two cancer related terms.

We investigated the occurrence of proteins and samples in biclusters belonging to one community (**Figure 1C**). A difference in the distributions of samples and features can be observed. The distribution of features is strongly positively skewed with a very low mode. In contrast, the sample distribution e.g. for communities 5 and 8 has a mode close to one. This shows that biclusters inside the same community (after filtering edges for random overlaps) can carry very different features and samples. By setting thresholds, ensemble biclusters can be restricted to point to consistent patterns in the data or to allow for variability.

In **Figure 1D**, we visualize the affiliation of biclusters to algorithms, which predicted them. This reveals that biclustering algorithms tend to identify overlapping regions in the data, resulting in highly connected communities consisting only of one algorithm. This shows the necessity of taking the results of multiple biclustering algorithms into account and not relying on one but many different algorithms to capture patterns in the data beyond the specificities of a single algorithm.

The results above demonstrate the power and utility of the workflow to establish a sophisticated biclustering analysis, to generate biological hypotheses. In the following we will examine MoSBi in a multi omics context.

### Individual biclustering algorithms vs. MoSBi

Next, we compared the individual performances of available biclustering algorithms and contrasted them with the performance of MoSBi. For that, we selected 6 published and publicly available datasets from the metabolomics, transcriptomics and proteomics disciplines (details in **Supplementary Table 1**). All datasets were analyzing cancer tissues or investigated cancer subtypes. As a gold standard, we used the condition match score to quantify the overlap between predicted biclusters and sample labels (see methods), where the relevance describes how well predicted biclusters correspond to known labels and recovery how well the labels were recovered by predictions. Additionally GO and KEGG pathway enrichment was performed to evaluate the gene sets in predicted biclusters.

The match between predictions and sample groups can be seen in **Figure 3A**. It reveals a heterogeneous performance of the individual biclustering algorithms. Spectral only predicted biclusters in two out of the six scenarios. Isa has the highest recovery on the Tang et al.^35^ metabolomics and Ku et al.^34^ proteomics data and both transcriptomics datasets, but has a poor performance on Yang et al.^36^ metabolomics and Wiśniewski et al.^37^ proteomics data. While having a good recovery, Isa never scores best on relevance. A similar behavior can be observed for Plaid, which on average performs very well for relevance, but shows low recoveries. It can also be observed that only plaid has in one proteomics dataset a relevance and recovery higher than 0.5. We then applied our ensemble approach on the predictions of all algorithms per dataset (**Figure 2A**, black marker). The ensemble approach is one of the two best performing tools in either recovery or relevance in all other datasets, except for the Tang et al.^35^ metabolomics data, where we could observe high overlaps with other clinical confounders (**Supplementary Figure 1**). On metabolomics data, with fewer features compared to sequencing data, the communities can additionally be visualized as co-occurrence networks (**Supplementary Figure 2**). Over all six datasets, MoSBi performed second best on average by Relevance and second best by recovery after plaid and Isa, which both have poorer performance on the other scale.

**Figure 2:**
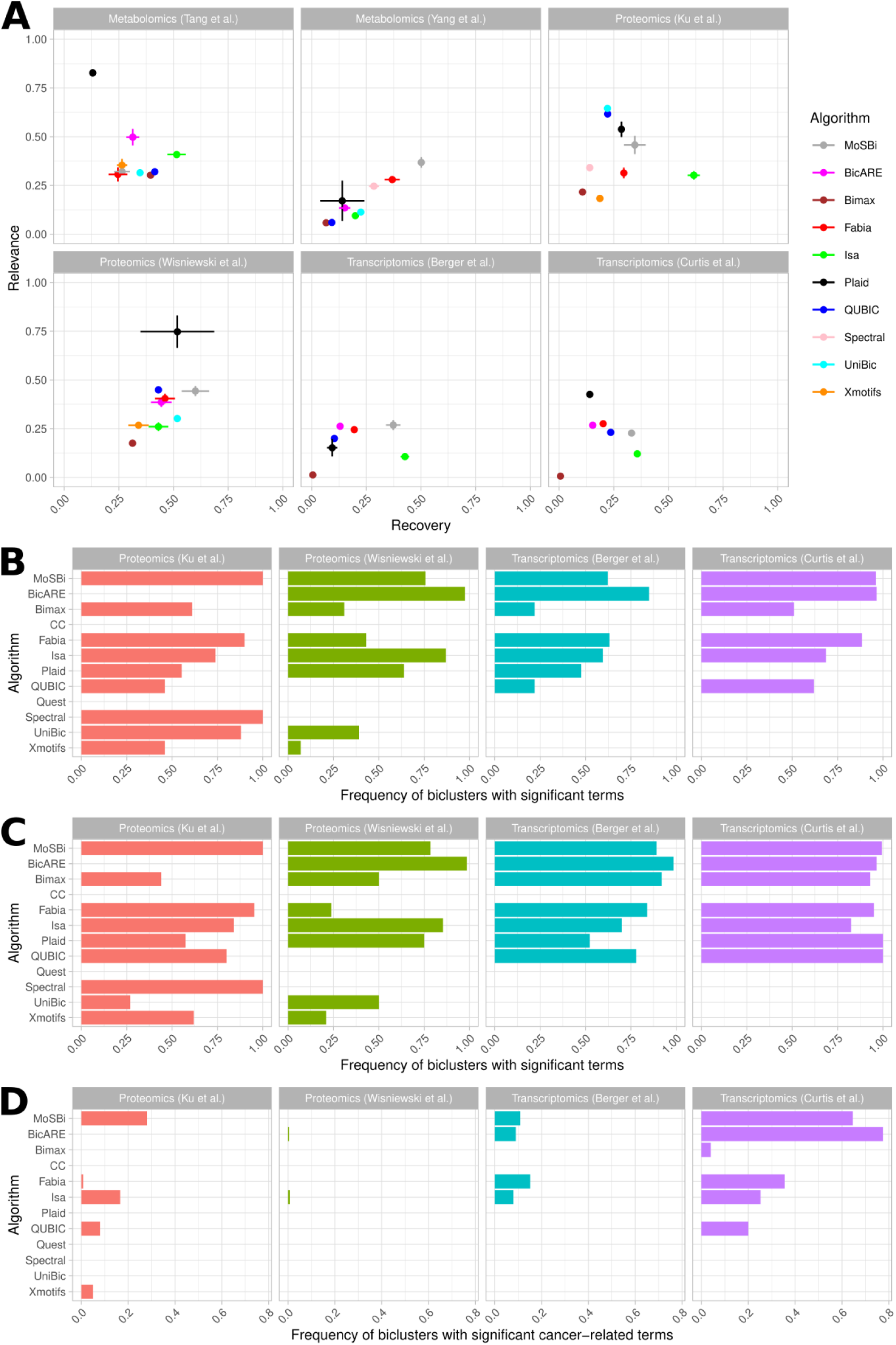
Performance of MoSBi and individual biclustering algorithms on cancer related omics data. For further information about the data see **Supplementary Table 1**. (**A**) Recovery and relevance for the condition match score of biclustering tools based on samples for cancer (subtypes). (**B**) Frequency of predicted biclusters per algorithm, with one or more significant KEGG terms (Adjusted p-value cut-off < 0.05). (**C**) Frequency of biclusters with one or more significant GO terms from the “biological process” category. (**D**) Frequency of predicted biclusters per algorithm, with one or more cancer related KEGG terms.

**Figure 3:**
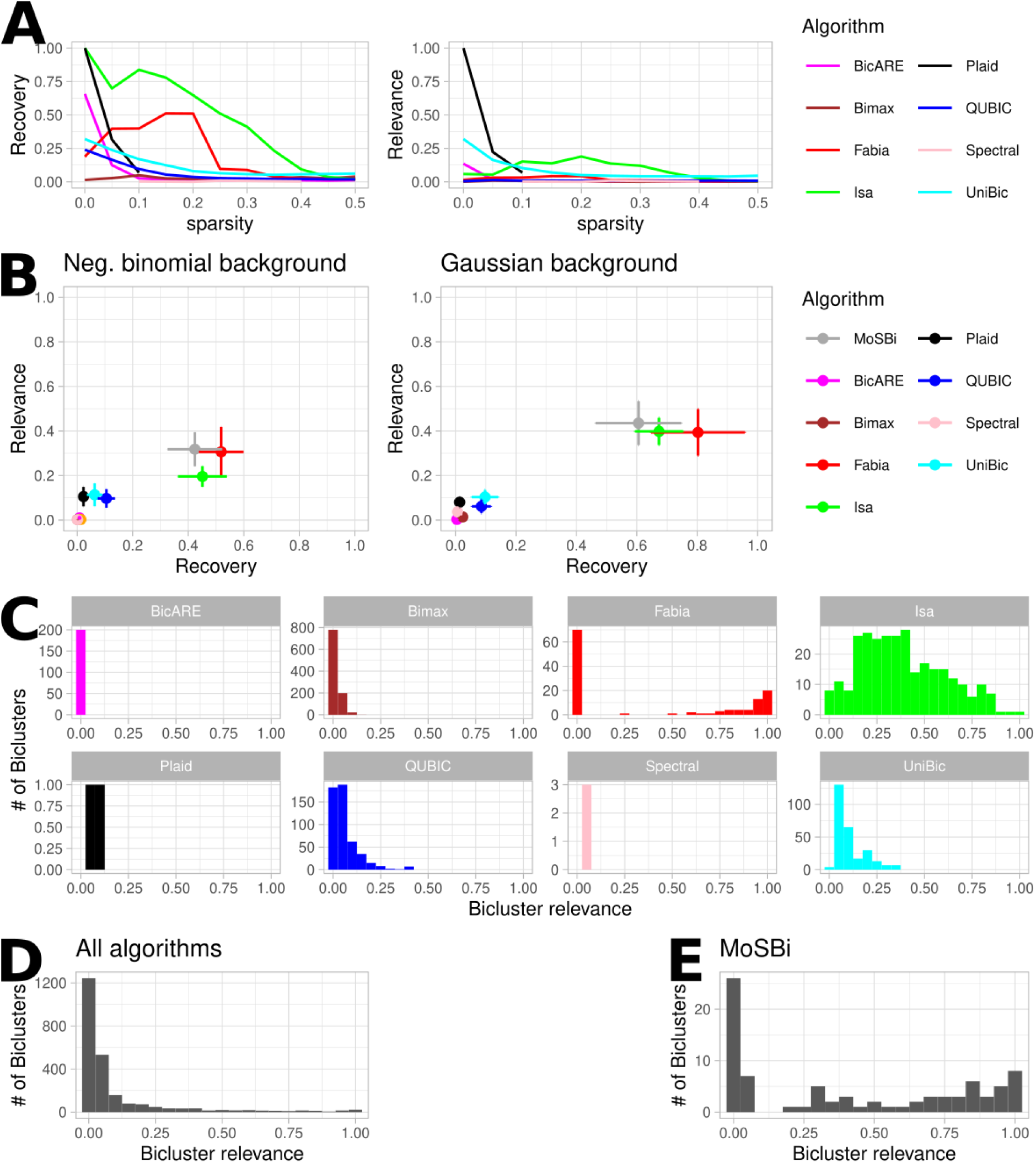
Evaluation of biclustering algorithms on synthetic data. (**A**) Recovery and relevance of biclustering algorithms with increasing sparsity, for one hidden shift bicluster. (**B**) Performance of biclustering algorithms and ensemble approach on a synthetic scenario including different bicluster types, sizes, sparsity and noise with a negative binomial distributed background (left) and normal distributed background (right). (**C**) Relevance distribution of biclustering algorithms for the scenario shown in (B, right). (**D**) Relevance distribution of all algorithms summed up from (C). (**E**) Relevance distribution of the predictions of the ensemble approach using the biclusters from (D).

To investigate the performance of the algorithms on the gene level, we performed KEGG pathway and GO biological process enrichment for the proteomics and transcriptomics datasets. In KEGG enrichment (**Figure 2B**), BicARE predicted the most biclusters with at least one significantly enriched term in three datasets. Interestingly, it did not stand out when investigating sample group labels. The ensemble method again showed a better performance than the average of biclustering algorithms. The same holds for enrichment of biological processes with GO terms (**Figure 2C**). Since all investigated proteomic and transcriptomic datasets were cancer related, we searched specifically for enriched KEGG pathways including the word “cancer”, “carcinoma” or “tumor” (**Figure 2D**). On the Wiśniewski et al.^37^ proteomics data, only Isa and BicARE found significant terms for biclusters, but only at very low frequencies. In the Ku et al.^34^ proteomics data, MoSBi found the most significant terms, on the two transcriptomics datasets, the second most after fabia and BicARE.

This reveals that individual biclustering algorithms peak in one or another measure or data set, but in an unpredictable manner. However, the MoSBi ensemble approach is more consistent and therefore more reliable for biclustering analysis.

### Performance on synthetic data

Evaluation on experimental data is preferable, since it accurately resembles real-life application of biclustering and stratification. Unfortunately, 2D gold standards are usually not available since many factors are influencing the molecular state of samples. Synthetic data can overcome this problem. This is frequently done to evaluate biclustering algorithms^6,25,38^.

Based on the synthetic data generation of Prelic et al.^25^, we developed a workflow to create synthetic scenarios, where one or multiple properties can be investigated (See **Supplementary Information**: Synthetic evaluation scenarios). We repeated previous scenarios from Prelic et al.^25^ and created novel scenarios, covering sparsity, overlaps and mixed sizes (**Supplementary Table 2**) and evaluated them on biclustering algorithms (See methods, **Supplementary Figures 5-9**). Since molecular omics data can include missing values, we investigated the effect of sparsity on the performance of biclustering algorithms (**Figure 3A**). While the overall performance of all algorithms decreases with increased sparsity, Fabia and Isa showed a higher resilience until a sparsity of 20% (percentage of missing values in the matrix, see **Supplementary Information**) after which the results deteriorated. The relevance was more robust against sparsity and did not decrease as strongly as the recovery.

So far, synthetic evaluation focussed on the assessment of individual characteristics of the data (e.g. noise or size). Using our workflow and knowledge from previous synthetic scenarios, we defined a complex scenario, incorporating all previously mentioned manipulations to the data (**Figure 3B**). We evaluated all approaches in this scenario and added a negative binomial background to simulate unique molecular identifier RNA sequencing data (**Figure 3B left**). Performance analysis separated the tools in two groups, clearly higher performing tools consisting of Fabia, Isa and MoSBi, and the rest performing significantly inferior. Fabia shows the best recovery and the ensemble approach the best relevance, however only marginally above Fabia and Isa. Even with a poor performance of many algorithms, it can still achieve high recovery and relevance. Algorithm selection has an influence on every ensemble approach, therefore, excluding the worst performing algorithms from the ensemble approach, yields a high increase of the relevance of the ensemble approach, while the recovery remains similar (**Supplementary Figure 10A**).

Being an average, the relevance does not characterize every distribution correctly, but is widely used in biclustering evaluation studies. We therefore investigated the relevance distribution of all algorithms independently (**Figure 3C**) and combined (**Figure 3D**). Some distributions are skewed. The combined distribution is positively skewed, showing that the majority of biclusters have very low overlap with the gold standard. Predictions by the ensemble approach show a different distribution (**Figure 3E**), where the majority of biclusters have a score above 0.5. Since an ensemble approach is sensitive to the performance of the underlying biclustering algorithms, we selected the best performing algorithms and repeated the analysis (**Supplementary Figure 10C&D**). As can be seen, the performance of MoSBi is even more evident, stating the importance of the utilized algorithms. By combining highly overlapping biclusters, MoSBi can reduce the number of mismatched biclusters. This also shows that the relevance distribution can give more detailed insights into algorithm performance. MoSBi additionally reduces the number of biclusters drastically, making an investigation of all predictions more manageable.

To investigate the performance of all algorithms under best conditions, we optimized their parameters to achieve the best possible performance (**Supplementary Figure 11**). This showed that algorithms can produce markedly better results given correct parameters compared to their standard parameters in **Figure 3B**. However this is time consuming and only possible for data with an existing gold standard. The differences between the two complex synthetic scenarios showed that parameters and performances vary widely between datasets. Therefore, an ensemble method offers an easier method to achieve good performance independently of parameter optimization.

### Biclustering in a multi-omics context

Since biclustering requires a data matrix as input, it can naturally be applied to multi-omics data, when combined to one data matrix. To investigate the performance of MoSBi in a multi-omics context, we used the TCGA breast cancer cohort from the Xena Platform^39^, which provides omics data for multiple breast cancer subtypes. RNAseq, miRNA and protein data was run independently and combined for all biclustering algorithms. All resulting bicluster networks (**Figure 4A**) appear similar, with big Basal communities and multiple communities consisting mainly of the LumA or LumB subtype, often highly interconnected. The protein data network shows a less distinct Basal community, whereas the miRNA data network shows Her2 samples mixed with LumB samples.

**Figure 4:**
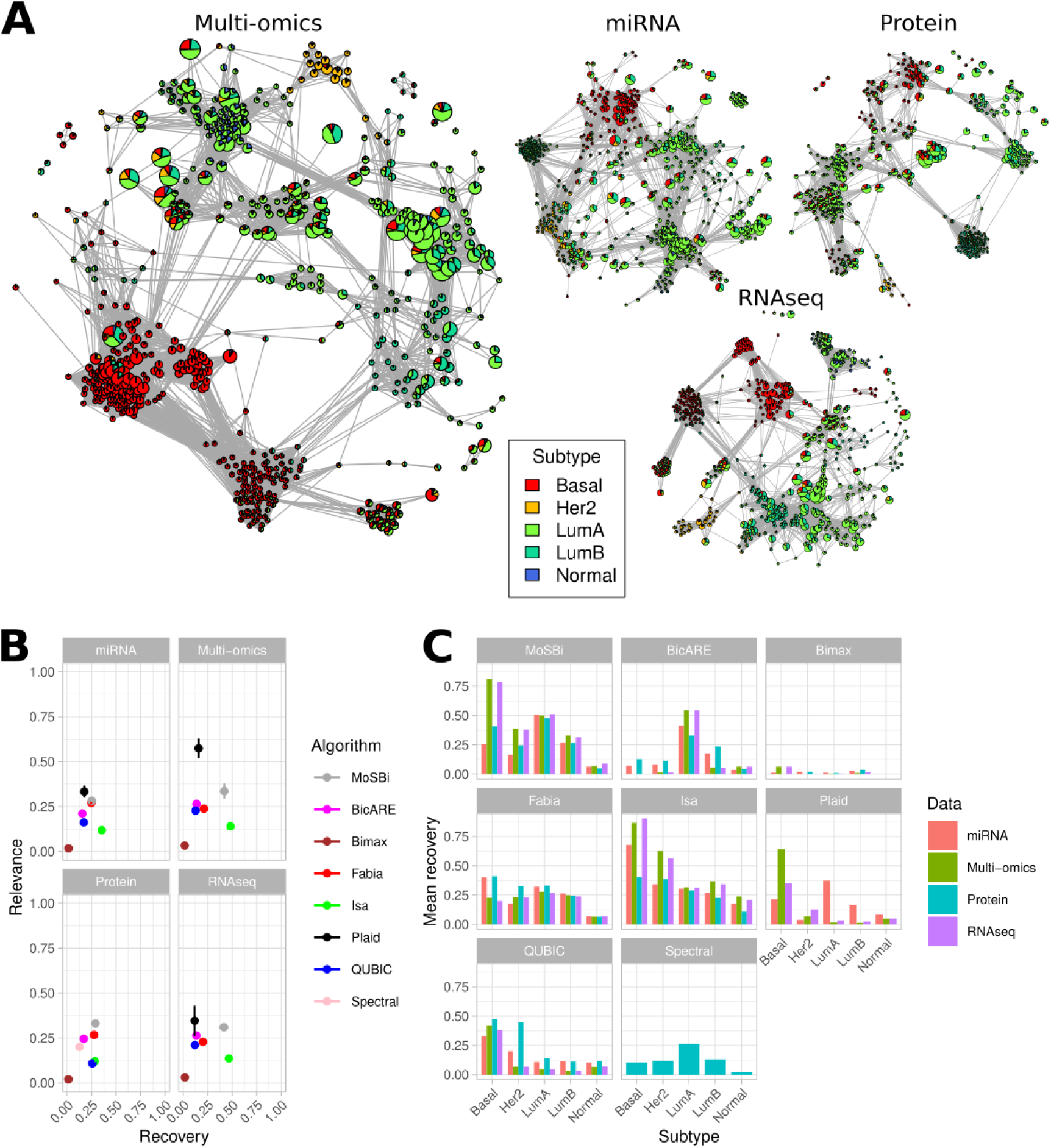
Biclustering on breast cancer multi-omics data. (**A**) Bicluster similarity networks on TCGA breast cancer miRNA, Protein expression, RNASeq, and combined data. Biclusters are colored by subtype and node size is proportional to sample size. (**B**) Relevance and recovery for the condition match score on the data sets from (A) for each algorithm individually and combined with MoSBi. All algorithms were executed 10 times. (**C**) Recovery for subtypes on the data sets from (A) for each algorithm individually and combined with MoSBi. All algorithms were executed 10 times.

In the next step, we evaluated the performance of the biclustering algorithms on the different data types. A consistent performance of most algorithms can be observed (**Figure 4B**), with only Plaid showing a high increase in relevance on the multi-omics data compared to the other data sets and did not identify any biclusters on the protein data. Only with MoSBi, a relevance and recovery higher than 0.25 could be observed in all four data sets. This shows that the multi-omics data did not yield a big performance increase for most algorithms, but rather that all data types carry the information to identify subtypes, with the ensemble approach being most robust throughout all data types.

While we did not find big differences on the overall performance, we next looked at the recovery of the subtypes individually (**Figure 4C**). Most algorithms did not recover all subtypes equally well. Isa has the highest recovery for Basal (above 0.75 for RNAseq and Multi-omics) and worst for Normal (All below 0.25). Fabia exhibits a more equal distribution, except for Normal, which has a low recovery throughout all algorithms. It can again be observed that all data types are similarly able to identify subtypes. In MoSBi, the Basal subtype has a better recovery in RNAseq and multi-omics data. Another interesting observation is that BicARE consistently recovers the LumA subtype through all data types.

In this analysis we can show that multi-omics biclustering is possible and can add value to the results. However, an individual biclustering analysis on all data types is also possible and yields similar performance. However, a combined analysis might be beneficial for a biological interpretation of biclusters, which consists of features from different omics types.

### The MoSBi software suite

To make our ensemble approach and biclustering algorithms in general, accessible for scientists and provide an easy to use interface, we developed the MoSBi suite for the Identification of *Molecular Signatures using Biclustering*. MoSBi is available as a R package (https://gitlab.lrz.de/lipitum-projects/mosbi) and web-app (https://exbio.wzw.tum.de/mosbi).

Many biclustering algorithms, such as Isa^28^ and Fabia^27^ or the biclust package use different result formats for returning biclusters. Therefore, we developed a unified framework, which is able to import predictions from various biclustering algorithms to simplify analysis of biclustering algorithms and apply our ensemble approach. Our network-based visualizations are also available in MoSBi which can be used with our ensemble approach or single biclustering algorithms. The framework can be extended to offer support for new biclustering algorithms and integrate them into the workflow. Networks can be exported as graphML for compatibility with tools such as Cytoscape^40^.

The web-app allows users without programming knowledge to stratify samples with our ensemble approach and profit from visualizations. Additionally, all biclustering algorithms can be accessed and executed with all parameters independently, if users are interested in specific algorithms. We also provide a docker image of the webtool, which allows the webtool to be deployed locally.

## Discussion

Stratification of patients based on molecular omics data is a challenging task and requires modern computational tools. Unsupervised approaches are suited to identify novel subgroups in the data. Biclustering is able to find meaningful patterns in modern omics data. In contrast to traditional clustering, algorithms output not only sample subgroups, but additionally feature subsets which characterize this similarity and can be further analyzed e.g. for functional associations to find disease mechanisms. We developed a biclustering ensemble approach, which takes the results of multiple biclustering algorithms and computes ensemble biclusters using a network-based approach. This is based on the assumption that biclustering algorithms predict highly overlapping biclusters, which we could show in our work. Various biclusters pointing to the same underlying data structures can indicate robust biclusters, which are then identified with MoSBi. We showed this on thymic epithelial tumor data^34^, where we were able to retrieve known cancer subtypes.

We demonstrated the application of MoSBi on cancer-related data sets and showed the possibility to perform a multi-omics analysis using biclustering. On various synthetic and experimental datasets, we assessed the performance of different biclustering algorithms and compared them to our ensemble approach. While Fabia and Isa on average performed best of all considered biclustering algorithms, no algorithm performed best in all scenarios and can be universally recommended. MoSBi did not always stand out but achieved a robust good performance in most scenarios. While the optimization of algorithm parameters on synthetic data could significantly improve the results, it leads to extensive run times and requires gold standard annotations, which are usually not available in real data, indicating that MoSBi is a preferable choice for biclustering. Additionally, it markedly reduces the number of biclusters. The network visualization gives an overview of the results and compared to other methods^17^ scales well with an increased number of biclusters.

The advantage of our ensemble approach over other biclustering ensemble approaches is that it is not algorithm-specific and via the MoSBi suite accessible as an API and graphical user interface. Unfortunately, some proposed approaches lack implementation^9,12^. Calculation of similarities between biclusters was proposed by Hanczar and Nadif^11^, where the authors calculated overlaps based on sums of overlaps of rows and columns, which can result in non-zero similarities for biclusters which share rows but no columns and therefore are in fact not overlapping (**Supplementary Figure 12**). They proposed their method as a single algorithm ensemble approach, did not filter for random overlaps and applied hierarchical clustering on the similarity matrix. This introduces another parameter for the number of consensus biclusters and assigns each bicluster to an ensemble bicluster, even with low overlap. Our approach avoids this by using the Louvain modularity. Additionally, MoSBi makes further analysis more accessible, since it markedly reduces the number of predictions, while not missing important biclusters. However, as an ensemble approach, MoSBi relies on the performance of multiple biclustering algorithms. We showed how the selection of biclustering algorithms can influence the results of MoSBi (**Supplementary Figure 10C&D**). While MoSBi is robust against a few badly performing algorithms, the majority of algorithms need to identify reasonable biclusters for MoSBi in order to work correctly. With new developments dn available algorithms, MoSBi can be extended to improve performance in the future

Our methodology offers a new perspective on biclustering and can visualize detailed properties of predictions. We demonstrated how a bicluster network analysis provides additional biological and structural insights into data. Clinical or experimental conditions can be associated with biological features. Using our approach, biclustering has the potential to play a significant role in disease subtyping and understanding.

## Methods

The biclustering ensemble algorithm consists of 4 major steps. These are execution of multiple biclustering algorithms, followed by a similarity computation for all returned biclusters, filtering of the similarity matrix for random overlaps and community detection on the similarity network. In the following all steps are described in detail:

### 1. Algorithms

Given an input matrix *M* ∈ ℝ^*R*×*C*^, we utilize different biclustering algorithms (**Table 1**) and collect their results in one combined list of biclusters *B* = [*B*_1_, *B*_2_, .., *B*_*n*_], where 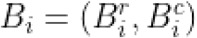 and 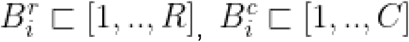 is a set of row and column indices of matrix *M* that belongs to bicluster *B*_*i*_. We implemented interfaces for all algorithms in our R package to generate this list using one unified API.

### 2. Similarity metrics

In the next step, pairwise similarities between all biclusters in *B* are computed. This is done using common similarity metrics, where the similarity is expressed as a two-dimensional overlap between biclusters. To do so, we treated a bicluster matrix as a two-dimensional area and computed their similarity in terms of overlapping areas. This is different to the additive similarity as proposed by Hanczar and Nadif^11^. One implemented metric is the Jaccard index. Our adaption resulted in the following formula:

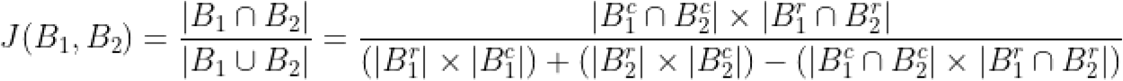

Besides the widely used Jaccard index, also the Bray-Curtis similarity, overlap coefficient and Fowlkes–Mallows index were implemented in a similar two-dimensional fashion. This results in a similarity Matrix *S* with *S*_*i,j*_ = *J*(*B*_*i*_, *B*_*j*_). Note that MoSBi can use any other definition of similarity as well.

### 3. Cut-off estimation

Since biclusters can have random overlaps that do not represent meaningful interactions, we estimate a cut-off to filter for such overlaps in the similarity matrix. This is done by randomly generating a list of biclusters *B*′ such that 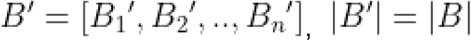 and 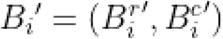 where 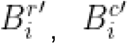 are randomly drawn without replacement from [1, …, *R*] and [1, …, *C*] correspondingly such that 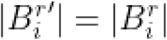 and 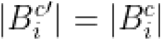.

To estimate the best cut-off *c*\* for the values in the similarity matrix *S*, we treat the *S* as an adjacency matrix and optimized for the biggest ratio between remaining edes in *S* and *S*′, where 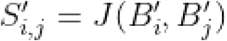 (To increase robustness, multiple randomizations *K* of *B* are used):

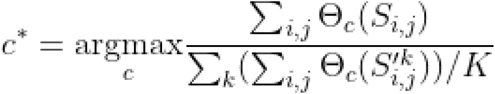with 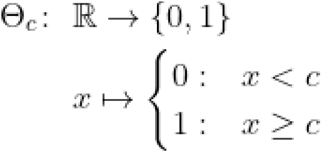. This results in the final and filtered similarity matrix *S*^*c*\*^where 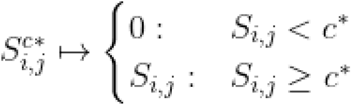.

### 4. Community detection

Finally, *S*^*c*\*^ is used as an adjacency matrix with biclusters as nodes and edges representing similarities. We compute the weighted Louvain modularity^41^, with similarities as weights, to find bicluster communities in the network. These highly similar bicluster communities can then be converted into ensemble biclusters using 3 parameters: `min_size` which defines the minimum number of biclusters in a community to convert a community into a bicluster, smaller communities are not considered. `row_threshold` & `col_threshold`, the minimum frequency of occurrence of a row/column element in a bicluster community to be taken over into an ensemble bicluster. E.g. with values of 0.5 only genes and samples will be part of the new ensemble bicluster if they occur in at least 50% of all biclusters in the corresponding community.

### Implementation

The workflow was implemented in the R programming language (version > = 3.6) and C++17. The web interface was realized with the Shiny web framework for R (version 1.4.0.2). The implementation for the R package is available here: https://gitlab.lrz.de/lipitum-projects/mosbi and for the web application here: https://gitlab.lrz.de/lipitum-projects/mosbi-webapp. The workflow can be executed from our web app on our servers or on a local machine using a public Docker image. For higher throughput or for the integration of our approach into a bioinformatics pipeline, the R package can be used directly.

### Visualizations

Network visualizations of the MoSBi package are implemented in R using the “igraph” package. Interactive plots in the webtool use the “visNetwork” library. All other visualizations use the “ggplot2” library in R.

### Co-occurrence Networks

For co-occurrence networks, biclusters from one community were selected. From this a new network is computed with samples and features as nodes. Edges can occur between samples and samples, samples and features and features and features. An edge is drawn between two nodes, if they occur together in at least one bicluster of the community. Edges are weighted by the number of biclusters, where two nodes co-occur. For the visualization, a network layout is computed, which takes the edge weights into account.

### Match score

The performance of biclustering algorithms and MoSBi was evaluated by comparing their overlap to labeled gold standard data. We used the commonly applied gene match score:

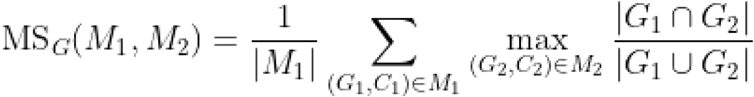

where *M*_1_ and *M*_2_ are two sets of biclusters, with each bicluster consisting of a set of genes *G*_*i*_ and conditions *C*_*i*_ (rows and columns)^25^. To investigate sample/condition overlaps, we define the according condition match score:

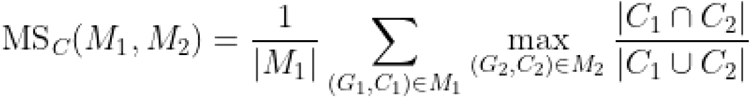

On synthetic data, where a two dimensional (2D) gold standard is available, we define the 2D match score as the multiplicative score of both dimensions:

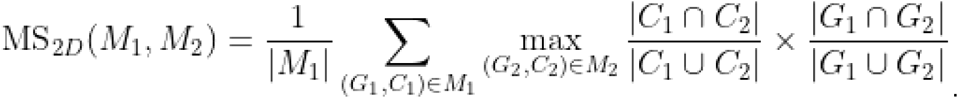

The scores can be used to compute relevance and recovery. Let *M*_*opt*_ be a set of implanted biclusters or a gold standard and *M* the output of a biclustering algorithm. Then, the average bicluster relevance is defined as MS(*M, M*_*opt*_) and describes to what extent the biclusters found by the algorithm correspond to the true hidden biclusters in the gene, condition or both dimensions. Similarly, the average bicluster recovery is defined as MS(*M*_*opt*,_ *M*) and describes how well each of the true biclusters is recovered by the algorithm. The recovery and relevance score both have an optimal value of 1 indicating a perfect overlap and 0 indicating no overlap.

The match scores describe a normalized sum of values. To investigate how well all individual biclusters predicted by one algorithm match the gold standard, we investigated the relevance distribution RD = [*rd*_1_, *rd*_2_,.., *rd*_*n*_] with *n* as the number of biclusters in set of biclusters *M* and

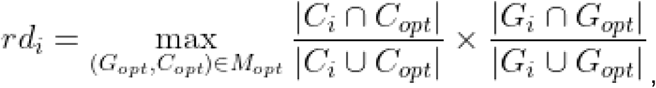

where *C*_*i*_ and *G*_*i*_ the columns and rows of bicluster *M*_*i*_.

### Experimental omics data

We evaluated the biclustering algorithms and MoSBi on six publicly available metabolomics (Tang et al.^35^, Yang et al.^36^), proteomics (Ku et al.^34^, Wiśniewski et al.^37^) and transcriptomics (Berger et al.^42^, Curtis et al.^43^) datasets (See **Supplementary Table 1**). Feature wise z-score were computed for all datasets, and prior to that log2 transformed (except for Ku et al.^34^ and Curtis et al.^43^, which already showed a normal distribution). Transcriptomics data was filtered for genes with 80% coverage in all samples and filtered 5000 most variant genes, to reduce algorithm runtime. Gene set/pathway enrichment was performed using the “clusterProfiler” R package using the “enrichGO” (biological process enrichment) and “enrichKEGG” functions.

TCGA Breast Cancer data was downloaded from the Xena Platform^39^. RNAseq transcriptomics data was processed as described above, miRNA and protein data was filtered for 80% coverage in all samples and z-score transformed. Only samples occurring in all three datasets were considered for the individual and multi-omics analysis, which resulted in 484 samples with measurements for all three data types.

### Synthetic data generation

To investigate the performance of tools in a controlled environment with a fully known gold standard, we developed a pipeline to generate synthetic datasets with implanted biclusters and additional properties such as noise and sparsity. The pipeline is shown in **Supplementary Figure 1**.

A detailed description of all synthetic scenarios is available in the supplementary information.

## Acknowledgments

TDR, NK, and JKP are funded by the Bavarian State Ministry of Science and the Arts in the framework of the Bavarian Research Institute for Digital Transformation (bidt, grant LipiTUM).

## Author contributions

TDR, JB, RR and JKP conceived and designed the study. TDR implemented the approach as a package. TDR and TB developed the webtool. TDR, KALL, TB, OC, FM, NK performed the evaluation of the algorithms and provided feedback about the implementation. All authors provided critical feedback about the study and helped in writing the manuscript.

