## Supplementary Information for "MoSBi: Automated signature mining for molecular stratification and subtyping"

---

### Supplementary Information

### Individual biclustering algorithms vs. MoSBi

**Supplementary Table 1: Datasets utilized for the comparison of individual biclustering algorithms vs. MoSBi**

| Publication | Omics type | # of groups | Data size (features x samples) |
|---|---|---|---|
| Wiśniewski et al.[2] | Proteomics | 3 | 10927 x 32 |
| Ku et al.[3] | Proteomics | 6 | 3209 x 90 |
| Berger et al.[4] | Transcriptomics | 5 | 5000 x 1063 |
| Curtis et al.[5] | Transcriptomics | 6 | 5000 x 1898 |
| Tang et al.[6] | Metabolomics | 6 | 399 x 30 |
| Yang et al.[7] | Metabolomics | 2 | 226 x 88 |

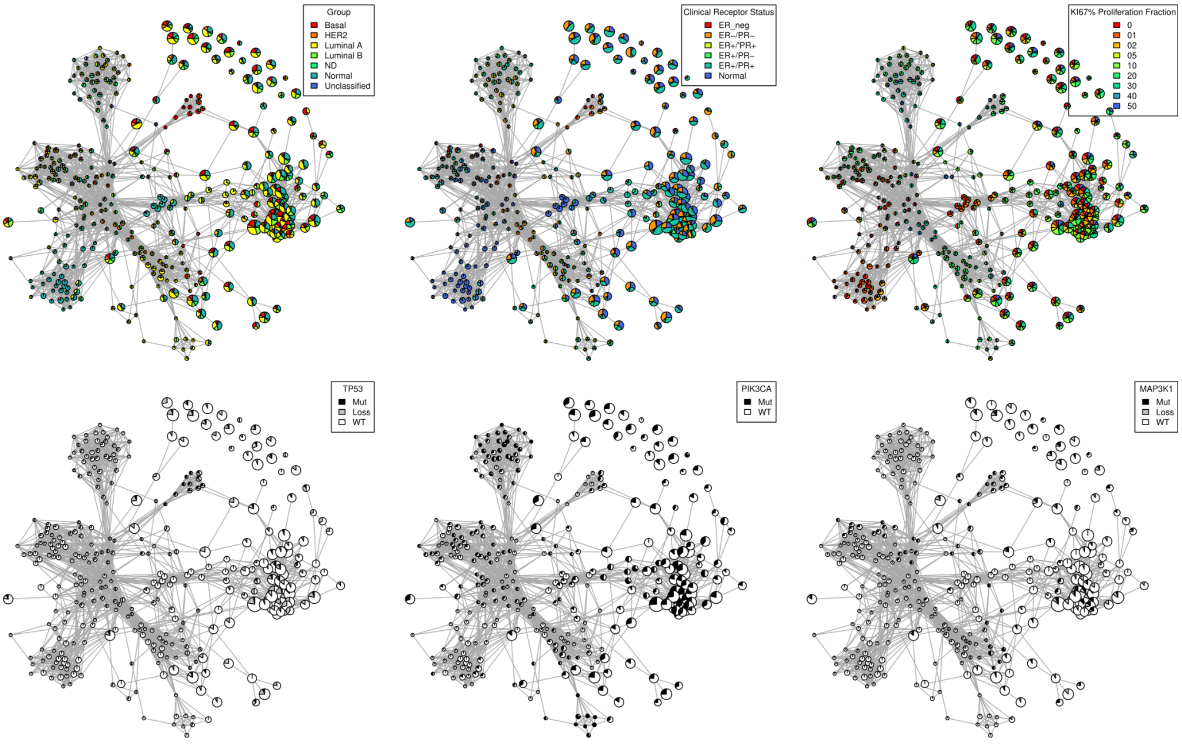


**Supplementary Figure 1: Bicluster networks of metabolomics data from Tang et al.[6]**
Biclusters are colored by clinical confounding factors, including cancer subtype, receptor status, KI67 proliferation and several mutations in known cancer driver genes.

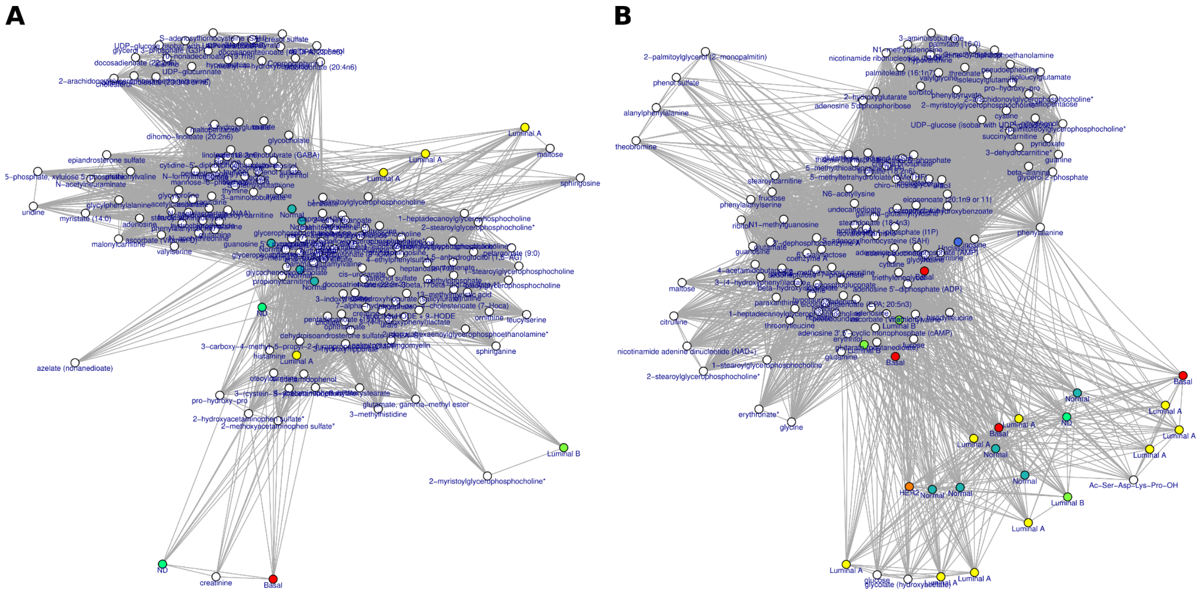


**Supplementary Figure 2: Co-occurrence networks of communities from Tang et al.[6]**
Exemplary Bicluster communities from the networks shown in Supplementary Figure 1. Nodes represent samples, colored by subtype and metabolic features (white). They are connected, if they co-occur in multiple biclusters of a Louvain community. For further details, see the Methods section.

### Synthetic evaluation scenarios

We first reproduced existing scenarios and took this as a starting point to develop further scenarios. We implemented and adapted the model described by Prelic et al.[8], focusing on the simultaneous row and column overlap. Additionally, we reproduced the dataset presented by Eren et al.[9], which used a bigger data matrix and multiple bicluster patterns. However, these benchmark datasets focused only on the classic scenarios like noise, overlap or bicluster pattern.

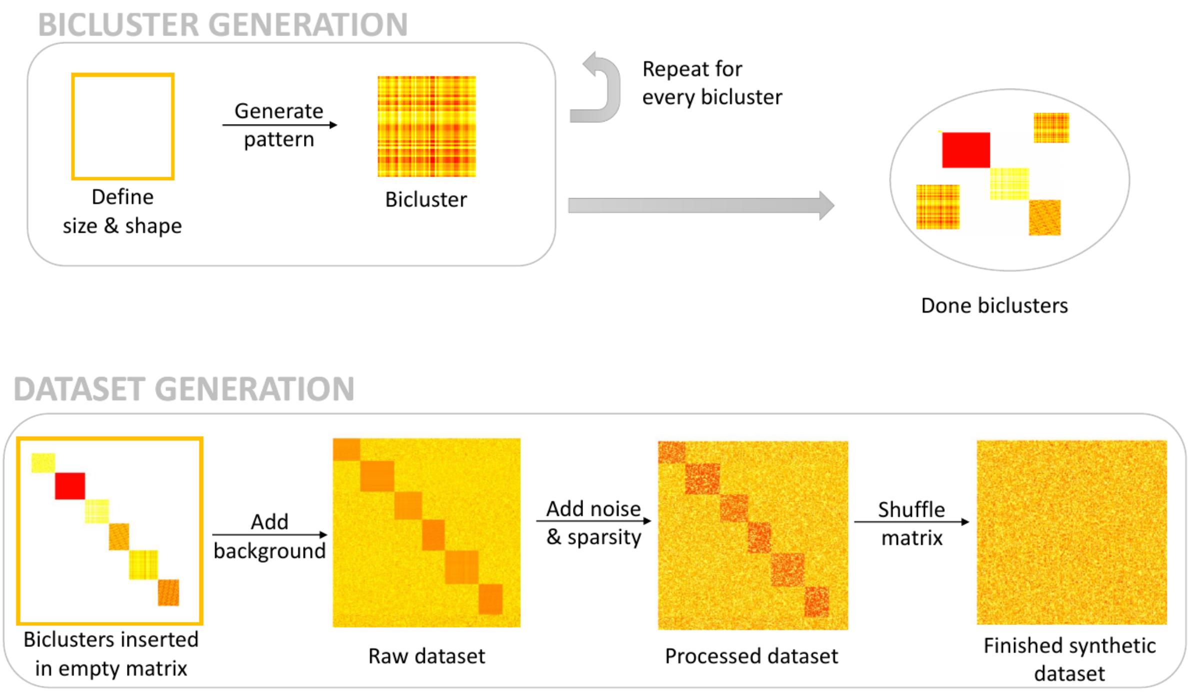


**Supplementary Figure 3: Data Generation Workflow.**
In the first phase, the biclusters are generated. After choosing the size and shape of the bicluster, the expression level is computed for a specific pattern. This process is repeated for every single bicluster. The second phase starts with an empty matrix, where the previously generated biclusters are inserted. The raw dataset is computed by adding a background distribution to the remaining empty cells. Next, the matrix is processed by adding noise and sparsity according to the previously set noise and sparsity levels. In the last step, the rows and columns of the matrix are being randomly permuted, resulting in the finished dataset.

The workflow of our data generation approach is illustrated in **Supplementary Figure 2**. First, biclusters with a previously defined shape, size and pattern are generated. The biclusters are then inserted in an empty data matrix and a background distribution is added to the remaining cells. The obtained raw matrix is processed by adding noise and sparsity. After performing a random permutation of rows and columns, the dataset is finished and ready to be used for the evaluation. Each data matrix has a corresponding gold standard, where the rows and columns of every hidden bicluster are listed. Before running the algorithms on the dataset, the matrix is normalized by computing the z-scores row-wise.

For the evaluation of biclustering algorithms, we generated a total of 670 synthetic datasets, grouped into 12 scenarios with 25 sub scenarios, covering many characteristics of real biological data. An overview of all datasets and their characteristics is shown in **Supplementary Table 1**. Each of them will be described in this section.

#### Pattern

All patterns are generated using a linear model. A bicluster $B = (I, J)$ of size $k \times l$ is defined according to the equation:

$$B = \begin{pmatrix} \pi_1\alpha_1+\beta_1 & \pi_2\alpha_1+\beta_1 & \cdots & \pi_l\alpha_1+\beta_1 \\ \pi_1\alpha_2+\beta_2 & \pi_2\alpha_2+\beta_2 & \cdots & \pi_l\alpha_2+\beta_2 \\ \vdots & \vdots & \ddots & \vdots \\ \pi_1\alpha_k+\beta_k & \pi_2\alpha_k+\beta_k & \cdots & \pi_l\alpha_k+\beta_k \end{pmatrix} \quad (1)$$

Where $\pi = \{\pi_1, \pi_2, \ldots, \pi_l\}$ is the base row vector, $\alpha = \{\alpha_1, \alpha_2, \ldots, \alpha_k\}$ is the scale column vector and $\beta = \{\beta_1, \beta_2, \ldots, \beta_k\}$ is the shift column vector. From that we define six bicluster patterns: constant, constant-row, constant-column, shift, scale, shift-scale[10]. In a constant bicluster, all elements have the same value over all rows and columns. This is illustrated in **Supplementary Figure 3A**. In a row-constant pattern, the elements located on the same row have equal values, but differ between distinct rows. An example of a row-constant pattern is shown in **Supplementary Figure 3B**. The opposite pattern is represented by the column-constant bicluster illustrated in Figure **Supplementary Figure 3C**. It has constant values on the column dimension and varying ones on the row dimension. The shift pattern is generated from a base vector $\pi$ and an additive component, represented by the shift vector $\beta$. This pattern is derived from the general linear model, with $\alpha = 1$ and shown in **Supplementary Figure 3D**. In a scale pattern, there is no additive component $\beta = 0$. The values inside the bicluster depend on the base row $\pi$ and the scale vector $\alpha$. An example is shown in **Supplementary Figure 3E.** The shift-scale pattern is the one corresponding to the general linear model, where the base row $\pi$, the scale column $\alpha$ and the shift column $\beta$ can take any value. **Supplementary Figure 3F** illustrates a shift-scale pattern.

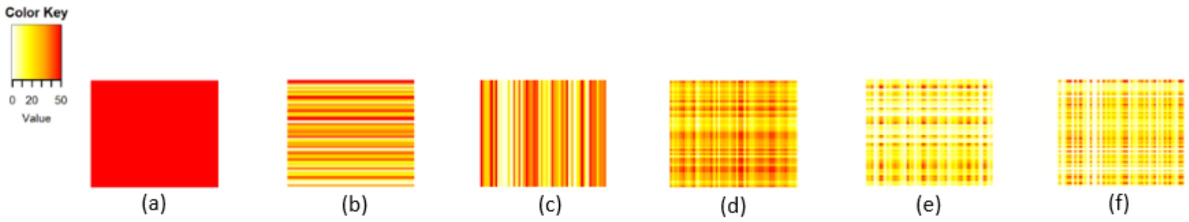


**Supplementary Figure 4: Examples of bicluster patterns.**
(a) Constant pattern, (b) row-constant pattern, (c) column-constant pattern, (d) shift pattern, (e) scale pattern, (f) shift-scale pattern. All patterns are particular cases of the general linear model presented in equation 1.

In a data matrix of size 500 x 200, one single bicluster is hidden. The bicluster of size 50 x 50 has one of the six patterns. The constant bicluster was generated with a constant expression value of 5, while the biclusters of other types were computed based on the previously defined linear model. The base, scale and/or shift vectors are randomly generated from U(1,50), where U(a,b) is the uniform distribution with a minimum of a and a maximum of b. Here, we distinguish two sub scenarios having different background distributions. The background matrix contains all cells of the matrix which do not belong to any bicluster. In the first scenario, the background matrix has an expression level of 0, while in the second sub scenario, the values in the background are randomly generated from U(1,100). Both sub scenarios contain 6 datasets each, one for every bicluster pattern.

#### Regulation

In this scenario we analyze the effect of different bicluster patterns when taking into account the regulation of the bicluster in relation to the background. An upregulated bicluster, which is higher expressed than the background distribution, is expected to be better detected by the algorithms than a bicluster whose expression values do not stand out from the background. We define the regulation of a bicluster in relation to the background matrix and distinguish 5 regulation levels of the bicluster: -2 (highly-downregulated), -1 (downregulated), 0 (no regulation), 1 (upregulated), 2 (highly-upregulated). First, we generate the bicluster expression levels in the same way we did for the scenario "Bicluster Pattern". Based on the bicluster values, the background distribution is created from a normal distribution with mean and standard deviation defined as it follows:

$$sd_{background} = sd_{bicluster} * U(1,3)$$
$$mean_{background} = mean_{bicluster} - (sd_{bicluster} + sd_{background}) * 2 * reg$$

Where $sd$ are the the standard deviations of the bicluster and background, respectively; $mean$ are the means of the bicluster and background; $reg \in \{-2,-1,0,1,2\}$ is the regulation factor and $U(a,b)$ the uniform distribution with values between a and b.

#### Noise

Studying the effect of noise on the performance of biclustering algorithms is one of the most frequently used scenarios when evaluating on synthetic data. The definition and implementation of noise might vary between publications. In this project, we define noise as a random number picked from a normal distribution with mean 0. The standard deviation represents the different noise levels: 0, 0.1, ..., 0.9, 1. For studying the effect of noise, we use a data matrix of size 500 x 200 and hide one highly-upregulated bicluster of size 50 x 50. The bicluster has a shift pattern which was computed using a base vector and a shift vector generated randomly from N(0,1). The background distribution is generated as described in the previous section. The bicluster is hidden in the data matrix and a z-score normalization is applied row-wise on the resulting matrix. After that, noise is added to every cell of the matrix. For a more robust result, the data is generated with 5 repetitions for each of the 11 noise levels.

#### Size and shape

Size and shape are other important bicluster characteristics, which may determine differences in the performance of biclustering algorithms. In these scenarios, investigate the effect of bicluster shapes and sizes on algorithm performances. For this reason, we use a 500 x 200 data matrix and a highly-upregulated shift bicluster of variable size and shape, generated as described before. For the "Bicluster Size" scenario, we use a square-shaped bicluster and test 11 different sizes: 5x5, 10x10, 20x20, ..., 90x90, 100x100. As a result, this scenario has 11 datasets. For the "Bicluster Shape" scenario, the lengths and the widths of the biclusters are limited to 6 different values: 30, 40, ..., 80.

#### Dataset size

In this scenario, we test whether the size of the dataset influences the biclustering result. Up to this point we used a data matrix with 500 rows and 200 columns. In this scenario, the dataset size is varied and takes each of the following sizes: 100x100, 200x100, 500x200, 1000x500, 5000x1000. The same shift bicluster of size 50 x 50 described previously is hidden in the data matrix.

#### Sparsity

Under sparsity we understand the percentage of values in a data matrix which are not available for technical or biological reasons (We set them to 0). For studying the effect of sparsity, we hide one shift bicluster of size 50 x 50 in a 500 x 200 data matrix. The shift pattern is computed using a base vector and a shift vector generated randomly from N(100,1). The background distribution is generated from a uniform distribution with the minimum value of 1 and the maximum value equal to the minimum value in the bicluster. In this way, we make sure that the bicluster is upregulated and all expression values in the matrix are greater than 0. We use 11 different sparsity levels: 0, 0.05, 0.1 ..., 0.45, 0.5. This means that 0 - 50 \% of the biclustering values are randomly set to 0.

#### Multiple Biclusters

All previously described scenarios involved hiding one single bicluster at a time. This scenario implants multiple biclusters of the same type in a data matrix. For this purpose, we use six variable biclusters of size 40 x 40 and a bigger dataset of size 500 x 300. The reason for the small change in size is the fact that we intend to place all six biclusters in the data matrix without any overlap. The previous scenarios revealed that these small changes in dimensions have no big effect on the performance of algorithms. All six biclusters are non-overlapping and upregulated. This scenario consists of three sub scenarios, each having one changed parameter. In the first simulation, we use biclusters of fixed pattern and size. All six biclusters have a shift pattern and the same size of 40 x 40. The pattern is generated with a base vector from N(100,20) and a shift vector from N(mean,20), where the mean is picked from U(30,100). In the second sub scenario, we vary the pattern of the biclusters and keep the size fixed. We use one bicluster of each pattern, all of them having size 40 x 40. For the third sub scenario, we use six shift biclusters with varying sizes of 30 x 50, 25 x 25, 40 x 60, 30 x 30, 45 x 45, 50 x 30.

#### One-dimensional overlap

Here we investigate the one-dimensional overlap of biclusters in a data matrix. The overlap can be either on the row dimension or on the column dimension. We define the overlap degree between two biclusters A and B as the number of rows/columns from bicluster A that are contained in bicluster B. The overlap simulation is done on 8 different overlap degrees: 0, 0.1, ..., 0.7, each with 10 repetitions. The analysis of one-dimensional overlap is repeated with row or column-overlap and biclusters of equal or different sizes.

#### Two-dimensional overlap

A two-dimensional overlap, i.e. simultaneous row and column overlap, defines a set of rows and a set of columns that belong to multiple biclusters at the same time. Generating this type of overlap with our own data generation approach is a difficult task, because expression levels that belong to two biclusters at the same time, should follow the pattern and fulfill the linear model of both biclusters. For this one scenario, we used a different approach and implemented the model introduced by Prelic et al.[8]. This consists of a 100 x 100 data matrix where multiple biclusters of the same size and pattern are hidden. In their publication, the authors describe two dataset types: constant and additive. The constant dataset is a binary data matrix, where the constant biclusters have an expression level of 1 and the background is 0. The additive dataset contains biclusters with column-constant patterns and the values in the background are generated from a uniform distribution, but are strictly smaller than the bicluster expression level.

#### Complex Scenarios

In these complex scenarios, we incorporated all aspects studied in the previous scenarios: multiple biclusters of different patterns, shapes and sizes, noise, sparsity and row- and column-overlap. For this simulation, a dataset of size 500 x 300, a noise level of 0.3 and a sparsity level of 0.1 were used. Six upregulated biclusters were placed in the data matrix. The pattern, size and shape of each bicluster were chosen randomly and independently for each dataset (which we repeated 10 times). It aims to reconstruct experimental omics data and is a difficult scenario, because of many inferring factors.

In addition, we generated three more scenarios following a slightly different approach. In these scenarios, we simulated the background distribution as a negative binomial distribution. In this way, the missing values in the dataset (simulated as sparsity) were already integrated in the distribution and did not have to be added synthetically. The sparsity level was varied by adjusting the parameters of the negative binomial distribution. This is meant to simulate unique molecular identifier RNA sequencing data[11].

**Supplementary Table 2: Synthetic data scenarios for evaluation on biclustering algorithms.**

| Scenario name | Subscenario Specification | Number of Biclusters | Bicluster Pattern | Bicluster Size | Regulation | Dataset Size | Noise | Sparsity | Overlap | Number of Datasets |
|---|---|---|---|---|---|---|---|---|---|---|
| **Bicluster Pattern** | No background | 1 | all | 50x50 | - | 500x200 | 0 | 0 | - | 6 |
| | Random background | 1 | all | 50x50 | - | 500x200 | 0 | 0 | - | 6 |
| **Bicluster Regulation** | Highly upregulation | 1 | all | 50x50 | 2 | 500x200 | 0 | 0 | - | 6 |
| | Upregulation | 1 | all | 50x50 | 1 | 500x200 | 0 | 0 | - | 6 |
| | No Regulation | 1 | all | 50x50 | 0 | 500x200 | 0 | 0 | - | 6 |
| | Downregulation | 1 | all | 50x50 | -1 | 500x200 | 0 | 0 | - | 6 |
| | Highly downregulation | 1 | all | 50x50 | -2 | 500x200 | 0 | 0 | - | 6 |
| **Bicluster Size** | Varying sizes | 1 | shift | 5x5<br>10x10<br>20x20<br>...<br>100x100 | 2 | 500x200 | 0 | 0 | - | 11 |
| **Bicluster Shape** | Varying shapes and sizes | 1 | shift | 30x30<br>30x40<br>...<br>30x80<br>40x30<br>40x40<br>...<br>40x80 | 2 | 500x200 | 0 | 0 | - | 36 |
| **Dataset Size** | Varying dataset sizes | 1 | shift | 50x50 | 2 | 100x100<br>200x100<br>500x200<br>1000x500<br>5000x1000 | 0 | 0 | - | 5 |
| **Noise** | Varying noise levels | 1 | shift | 50x50 | 2 | 500x200 | 0<br>0.1<br>...<br>1 | 0 | - | 55 |
| **Sparsity** | Varying sparsity levels | 1 | shift | 50x50 | 2 | 500x200 | 0 | 0<br>0.05<br>...<br>0.5 | - | 55 |
| **Multiple Biclusters** | Same pattern | 6 | shift | 40x40 | 1 | 500x300 | 0 | 0 | - | 5 |
| | Varying patterns | 6 | all | 40x40 | 1 | 500x300 | 0 | 0 | - | 5 |
| | Different sizes | 6 | shift | 30x50<br>25x25<br>40x60<br>30x30<br>45x45<br>50x30 | 1 | 500x300 | 0 | 0 | - | 5 |
| **One-dimensional Overlap** | Column overlap | 6 | shift | 40x40 | 1 | 500x300 | 0 | 0 | column<br>0<br>0.1<br>...<br>0.7 | 80 |
| | Row overlap | 6 | shift | 40x40 | 1 | 500x300 | 0 | 0 | row<br>0<br>0.1<br>...<br>0.7 | 80 |
| | Column overlap, varying bicluster sizes | 6 | shift | 30x50<br>25x25<br>40x60<br>30x30<br>45x45<br>50x30 | 1 | 500x300 | 0 | 0 | 0<br>0.1<br>...<br>0.7 | 80 |

| Scenario name | Subscenario Specification | Number of Biclusters | Bicluster Pattern | Bicluster Size | Regulation | Dataset Size | Noise | Sparsity | Overlap | Number of Datasets |
|---|---|---|---|---|---|---|---|---|---|---|
| | Row overlap, varying bicluster sizes | 6 | shift | 30x50<br>25x25<br>40x60<br>30x30<br>45x45<br>50x30 | 1 | 500x300 | 0 | 0 | 0<br>0.1<br>...<br>0.7 | 80 |
| **"Close to Real"** | Complex | 6 | random | random | 1 | 500x300 | 0.3 | 0.1 | random | 10 |
| **Two-dimensional Overlap)** | Constant model (binary matrix) | 10 | constant | $\sim$10x10 | - | $\sim$100x100 | 0 | 0 | row and column 0 - 0.8 | 45 |
| | Additive model | | column-constant | $\sim$10x10 | - | $\sim$100x100 | 0 | 0 | row and column 0 - 0.8 | 45 |
| **Negative Binomial Background)** | Sparsity | 1 | shift | 50x50 | - | 500x200 | 0.3 | 0 - 0.5 | - | 11 |
| | Complex without noise | 6 | random | random | - | 500x300 | 0 | $\sim$ 0.1 | random | 10 |
| | Complex without noise | 6 | random | random | - | 500x300 | 0.3 | $\sim$ 0.1 | random | 10 |

### Evaluation of biclustering algorithms on synthetic scenarios

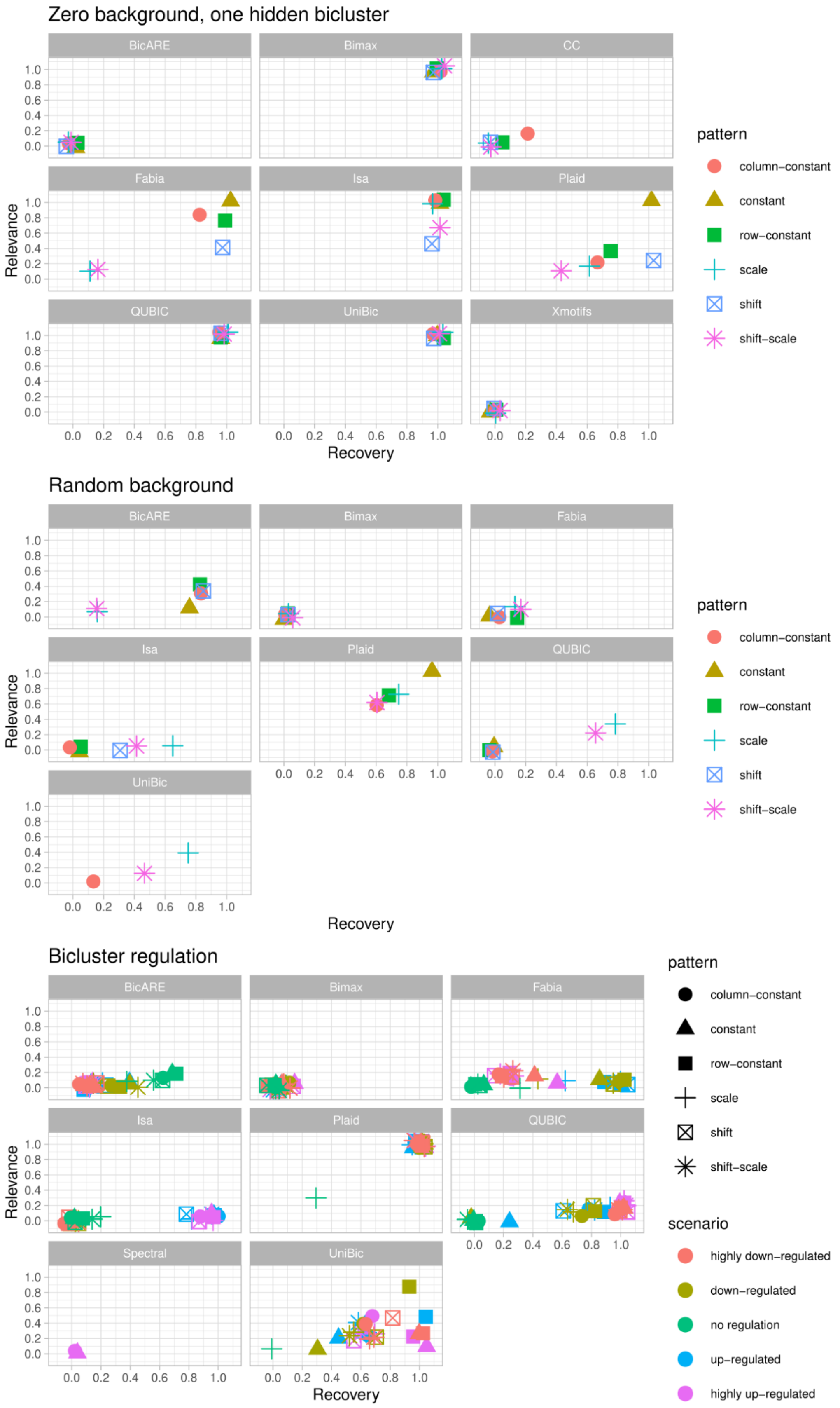


**Supplementary Figure 5: Evaluation on synthetic data.**

Datasets with a single hidden bicluster per data with no background, normally distributed background, biclusters of different factors.

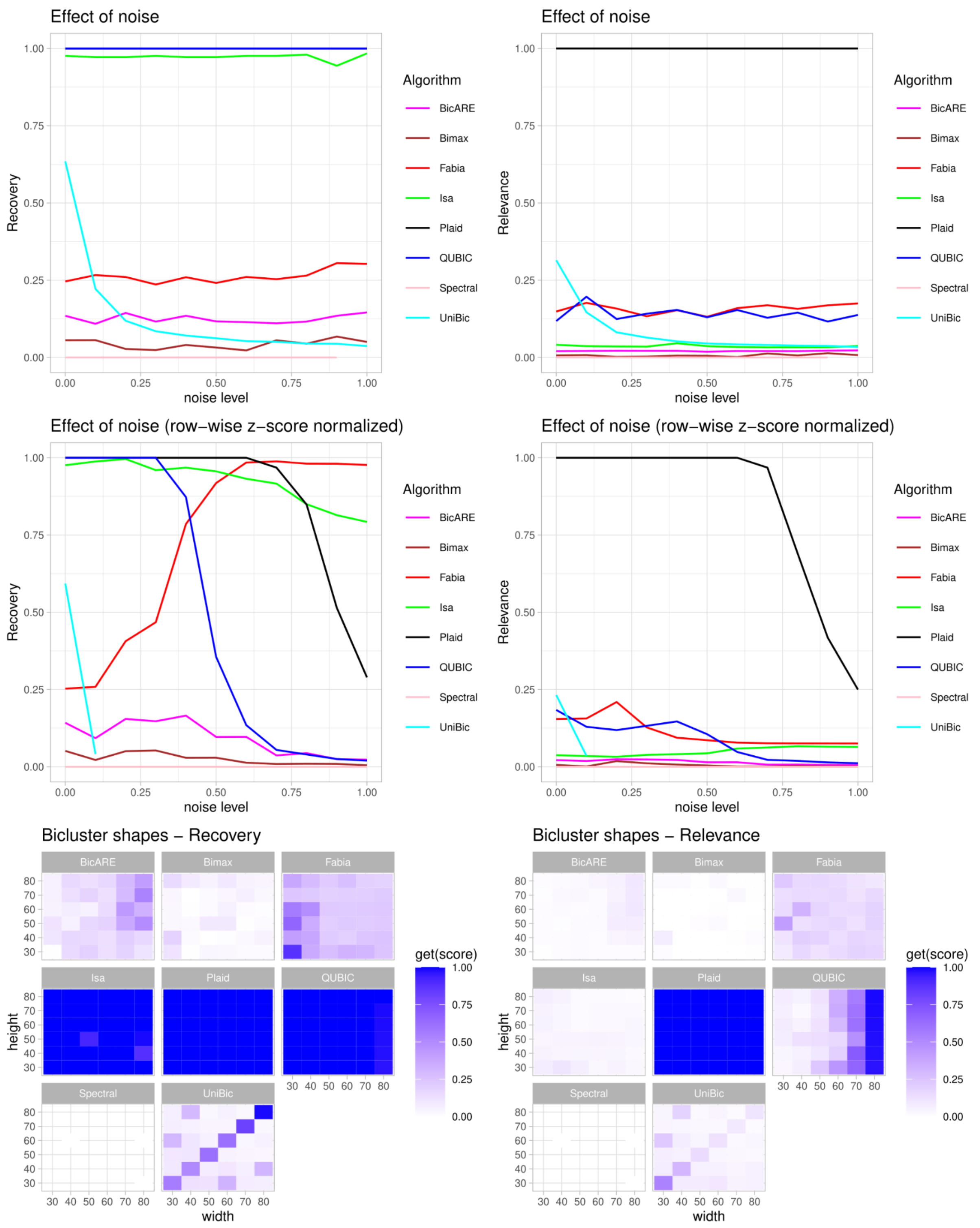


**Supplementary Figure 6: Evaluation on synthetic data.**
Performance on increasing noise levels with and without z-score normalization of the datamatrix. Performance on datasets with biclusters of different shapes.

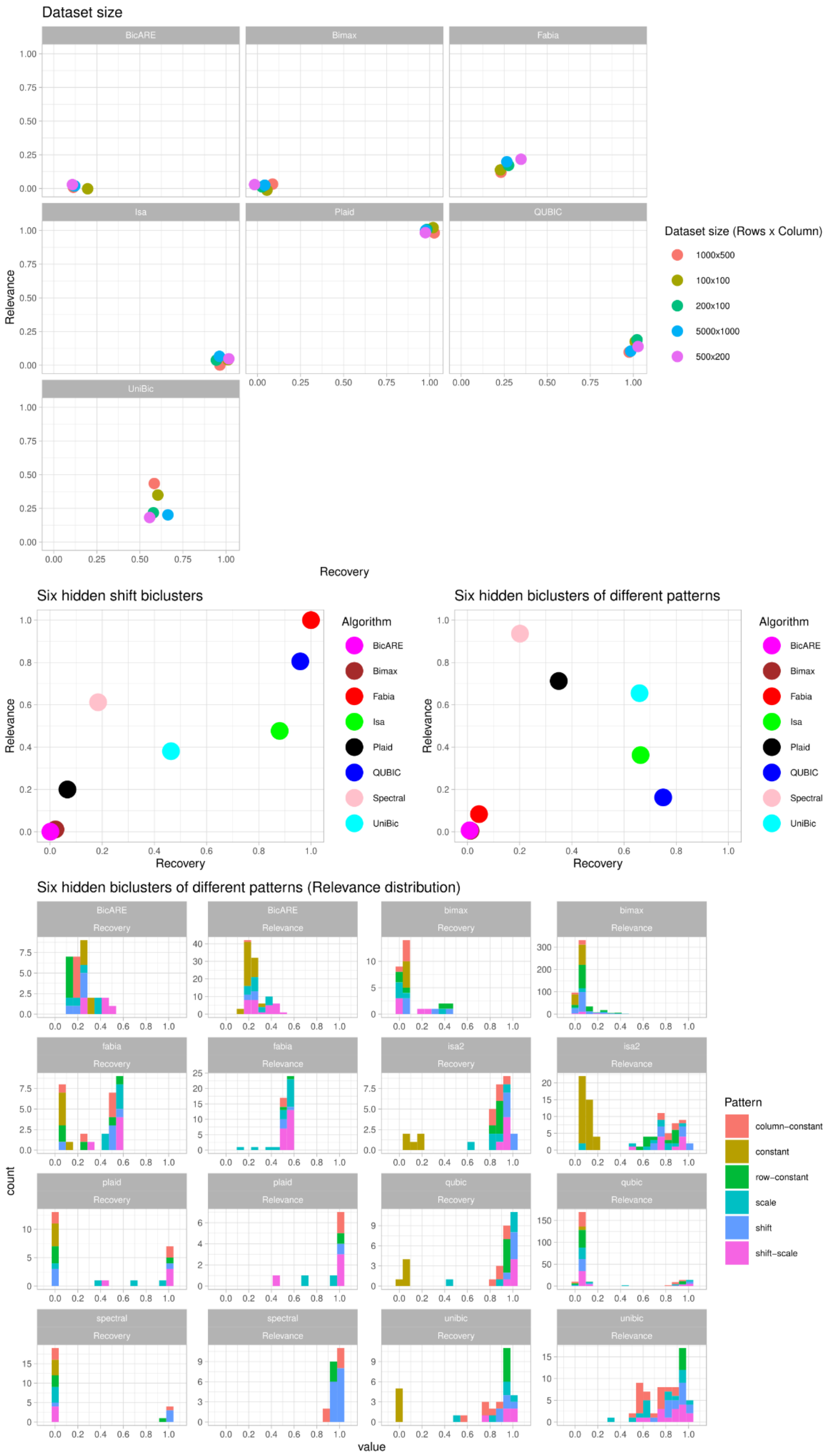


**Supplementary Figure 7: Evaluation on synthetic data.**
Top: Performance on varying dataset sizes. Middle: Six biclusters (of different patterns) hidden in the same matrix. Bottom: Relevance distribution six biclusters of different patterns hidden in the same matrix.

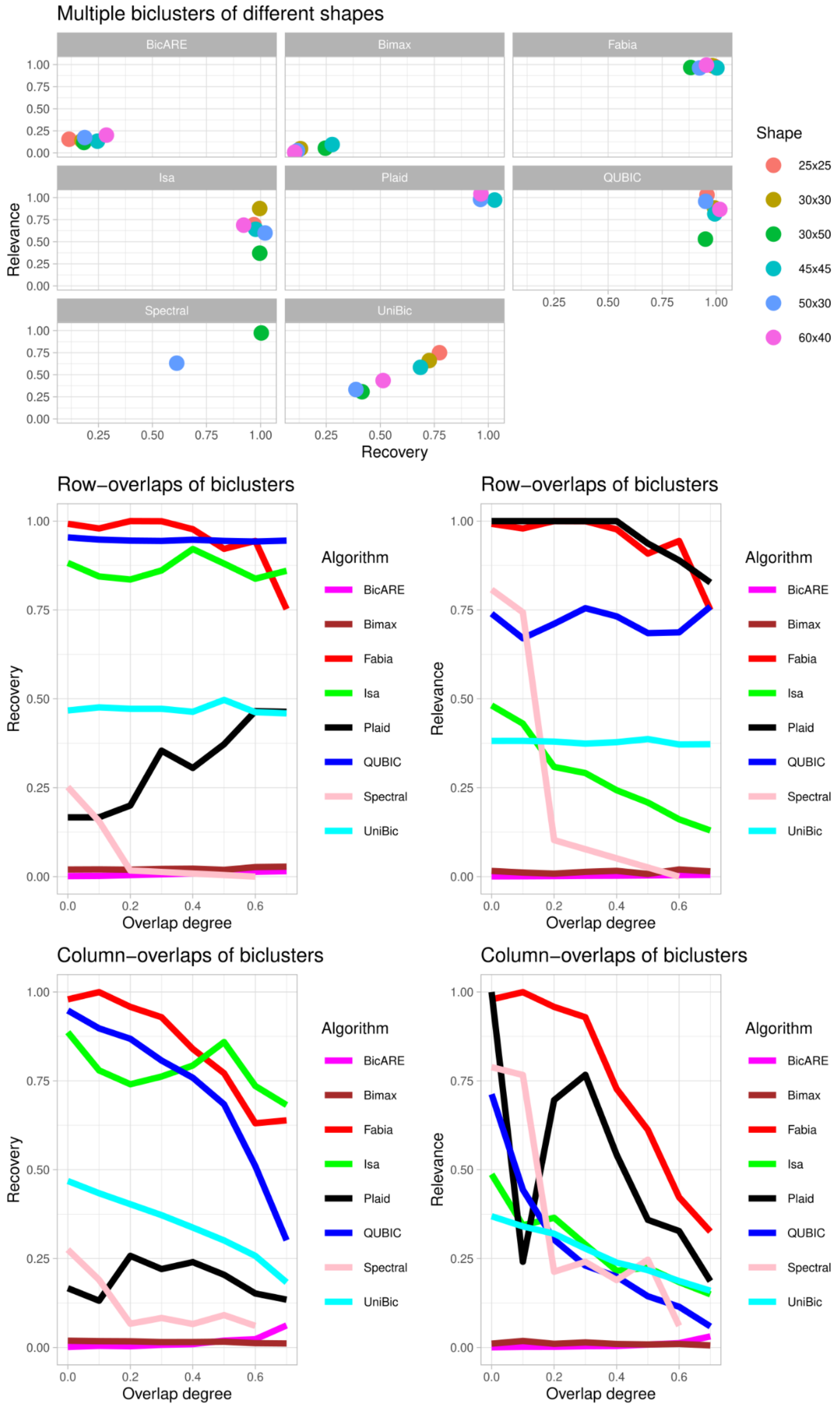


**Supplementary Figure 8: Evaluation on synthetic data.**
Top: Multiple biclusters of different shapes hidden in the same matrix. Middle: Multiple biclusters with overlapping rows in the same matrix. Bottom: Multiple biclusters with overlapping columns in the same matrix.

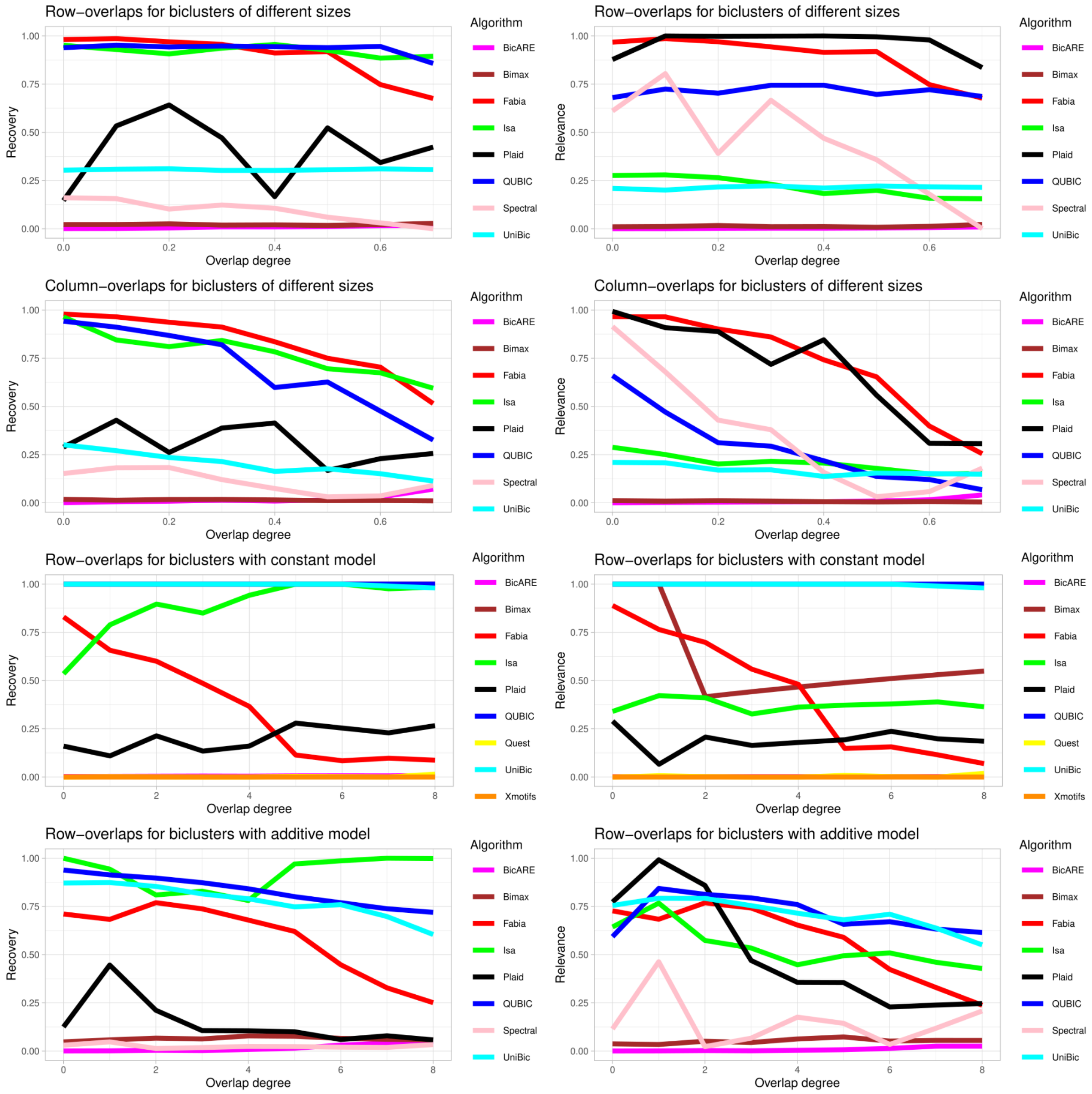


**Supplementary Figure 9: Evaluation on synthetic data.**
Multiple biclusters with overlapping rows or columns in the same matrix using a constant or additive model.

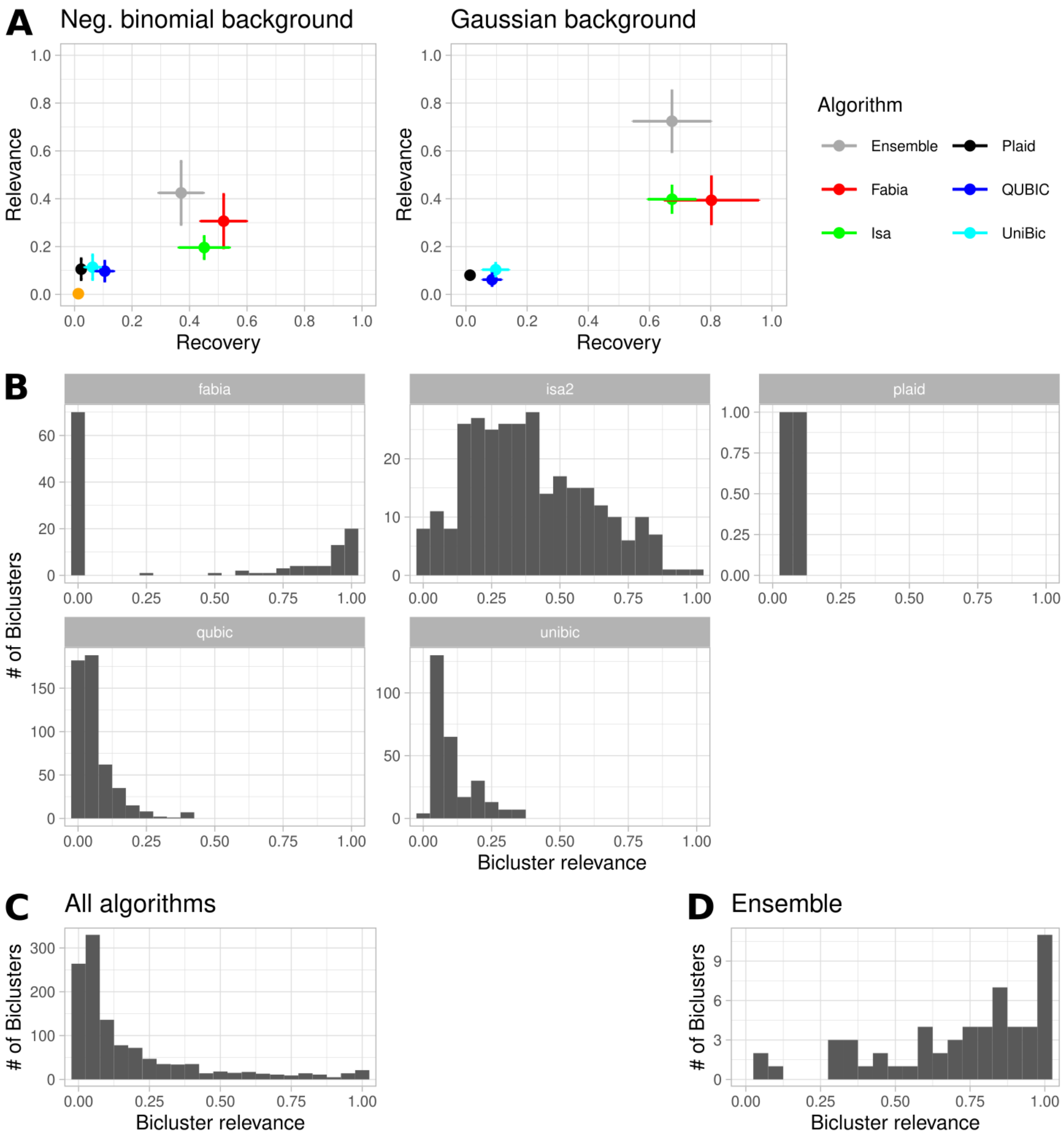


**Supplementary Figure 10: Synthetic evaluation with filtered algorithms.** For this figure only the biclustering algorithms Plaid, Fabia, QUBIC, Isa and UniBic were utilized. (A) Performance of biclustering algorithms and ensemble approach on a synthetic scenario including different bicluster types, sizes, sparsity and noise with a negative binomial distributed background (left) and normal distributed background (right). (B) Relevance distribution of biclustering algorithms for the scenario shown in (A, right). (C) Relevance distribution of all algorithms summed up from (B). (D) Relevance distribution of the predictions of the ensemble approach using the biclusters from (D).

### Parameter optimization for biclustering algorithms

We optimized the parameters for the biclustering algorithms using naive grid optimization as performed in the clustering methods evaluation by Wiwi et al.[12]. Where constraints for parameters were available by the tool's documentation, these were used otherwise defined by us. Each tool was given 1000 runs. The performances of all runs for all tools is plotted in **Supplementary Figure 8**.

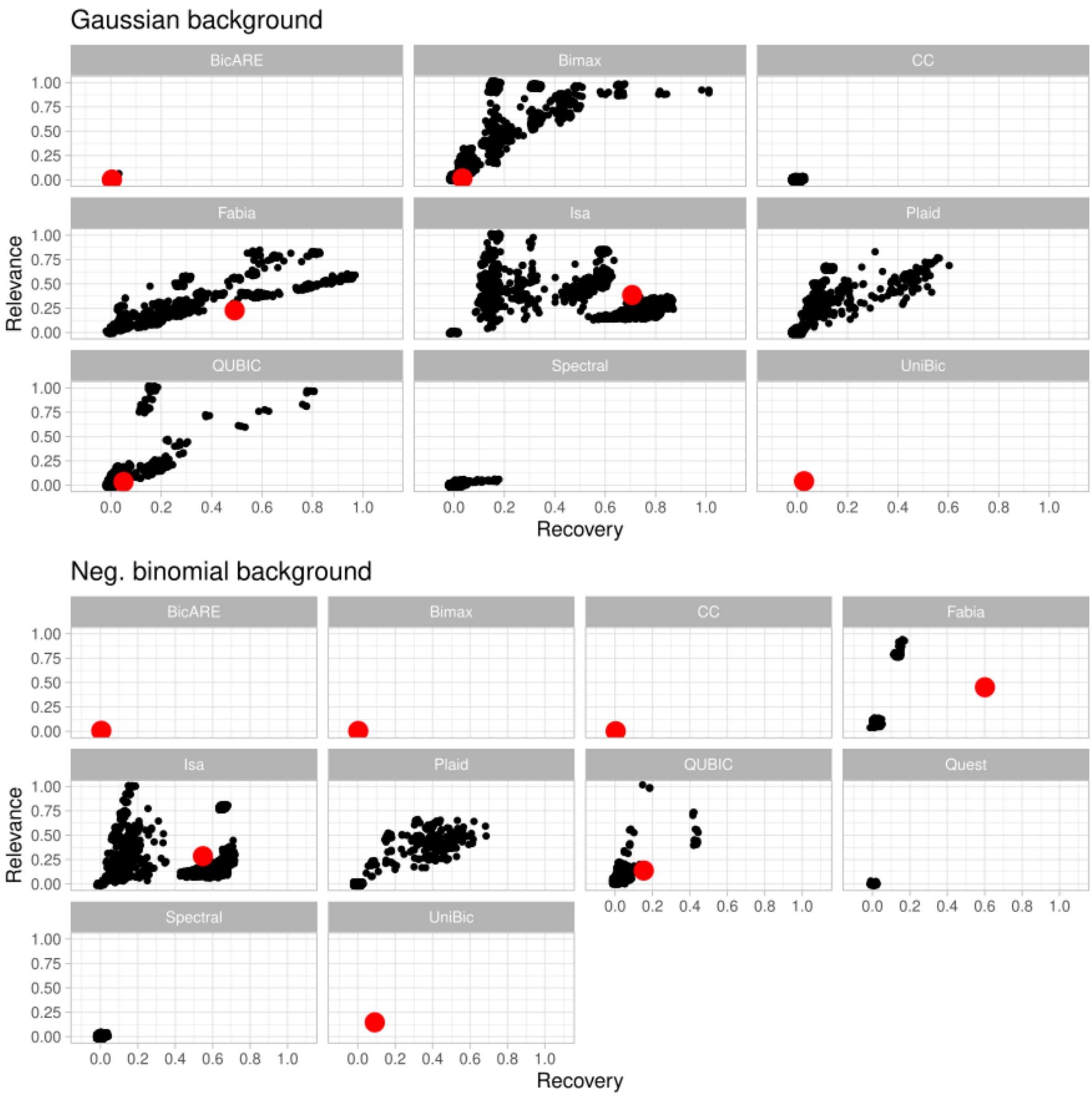


**Supplementary Figure 11: Optimization of parameters**
Optimization of parameters for biclustering algorithms on the two “complex” synthetic scenarios with a gaussian or negative binomial background distribution. The red dot showsicates the performance on default parameters.

### Bicluster similarity

Comparison of additive and multiplicative operlaps of biclusters. An additive score considers a one dimensional overlap as a non-zero overlap, while a multiplicative score only returns non-zero similarities for two-dimensional overlaps (**Supplementary Figure 11**).

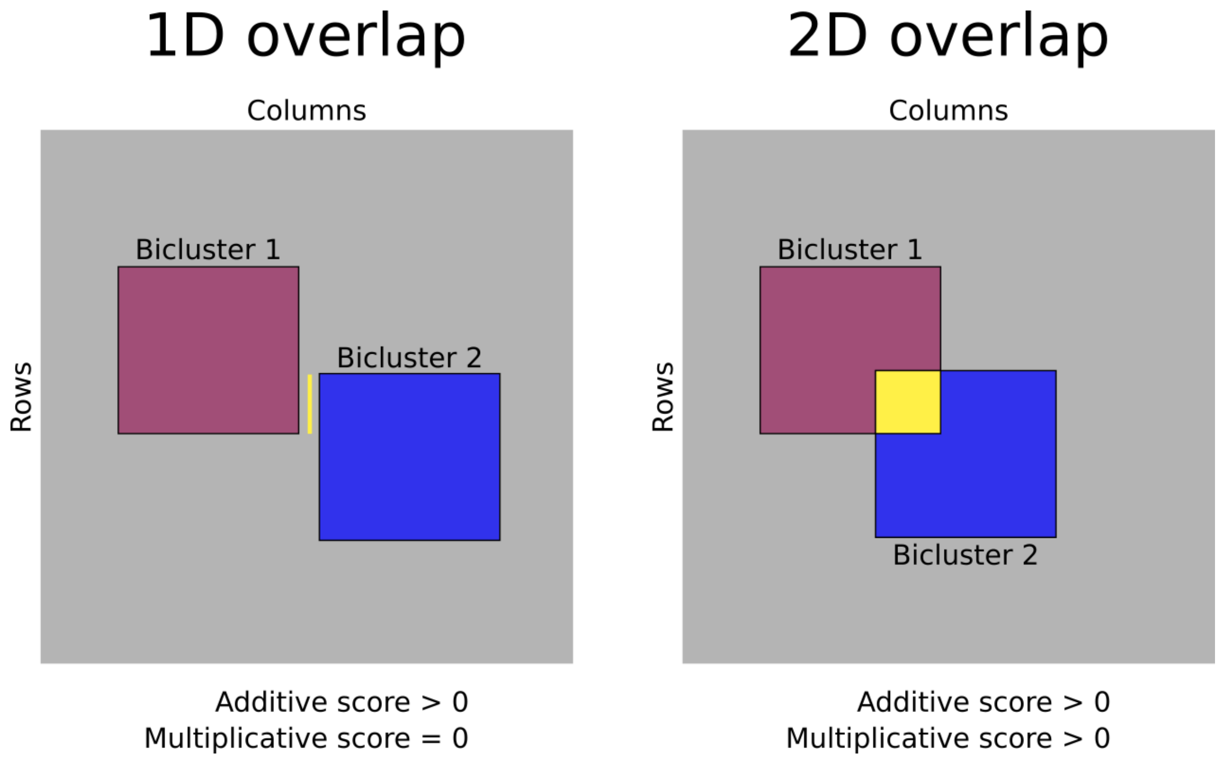


**Supplementary Figure 12: Comparison of additive and multiplicative overlaps of biclusters.**

Additive score as used by Hanczar and Nadif[13] to evaluate bicluster similarity and multiplicative score as used in this manuscript (See methods section) for one and two dimensional overlaps of biclusters.
